# No ‘small genome attraction’ artifact: A response to Harish et al. ‘Did viruses evolve as a distinct supergroup from common ancestors of cells?’

**DOI:** 10.1101/050179

**Authors:** Arshan Nasir, Kyung Mo Kim, Gustavo Caetano-Anollés

## Abstract

In a recent eLetter and associated preprint, Harish, Abroi, Gough and Kurland criticized our structural phylogenomic methods, which support the early cellular origin of viruses. Their claims include the argument that the rooting of our trees is artifactual and distorted by small genome (proteome) size. Here we uncover their aprioristic reasoning, which mingles with misunderstandings and misinterpretations of cladistic methodology. To demonstrate, we labeled the phylogenetic positions of the smallest proteomes in our phylogenetic trees and confirm that the smallest genomes were neither attracted towards the root nor caused any distortions in the four-supergroup tree of life. Their results therefore stem from confusing outgroups with ancestors and handpicking problematic taxa to distort tree reconstruction. In doing so, they ignored the details of our rooting method, taxa sampling rationale, the plethora of evidence given in our study supporting the ancient origin of the viral supergroup and also recent literature on viral evolution. Indeed, our tree of life uncovered many viral monophyletic groups consistent with ICTV classifications and showed remarkable evolutionary tracings of virion morphotypes onto a revealing tree topology.

---

Harish et al. would like to see the origin of Eukarya at the base of the Tree of Life (ToL) [1]. So, in their commentary [2,3], they begin by questioning our phylogenomic analysis, which is supported by large-scale structural and functional data and well-established comparative genomics, phylogenomics, and multidimensional scaling approaches [4]. Their writings [2,3] fail to acknowledge recent literature, including our recent invalidation of both their rationale [5] and their own phylogenetic methodologies [6], which we showed counter modern evolutionary thinking. While our critique remains unanswered, here we recap how their argumentation and misguided experimentation is used to *“mud the waters”* (*sensu* [2]) of cladistics understanding. A quick fact-checking exercise is presented in Table 1 and described in the following sections.

**Table 1.**
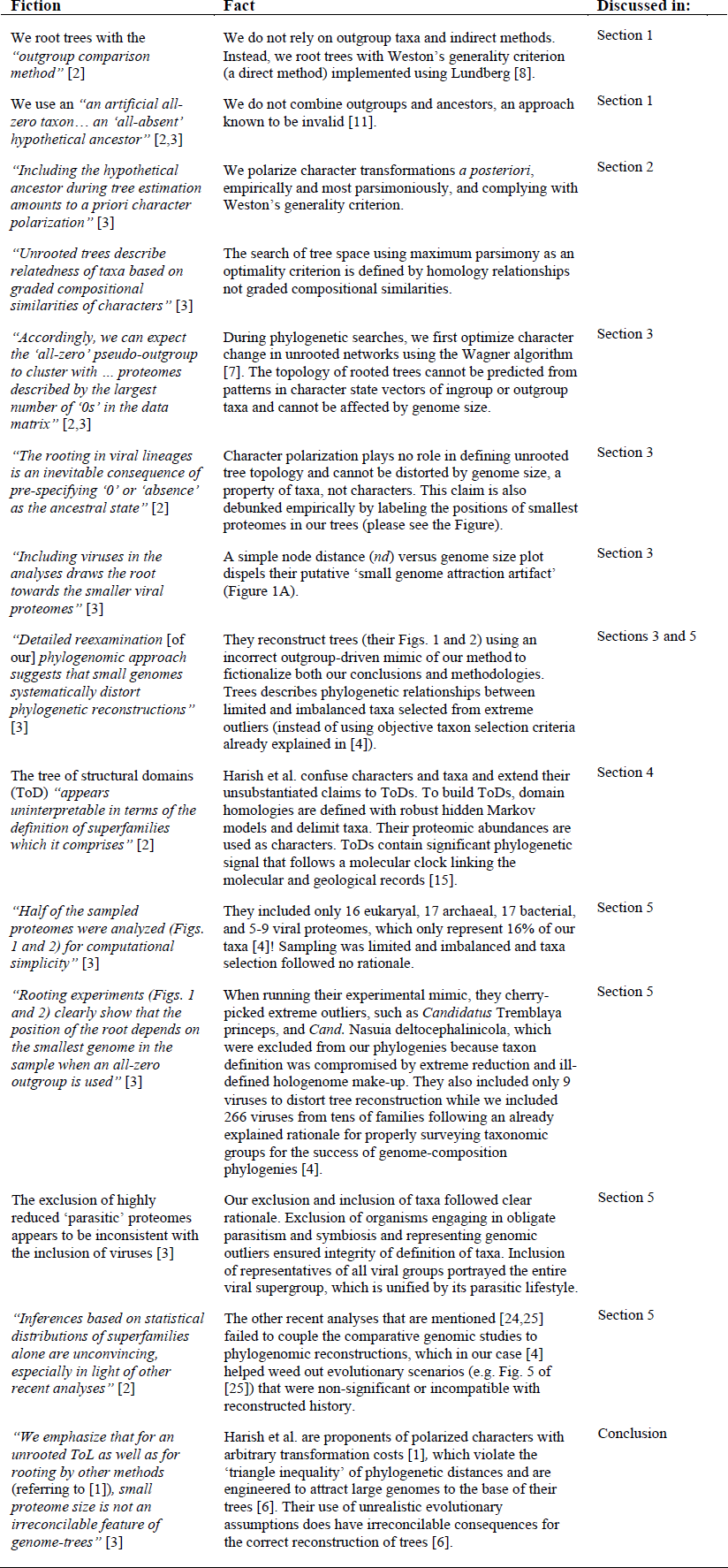
Fact-checking the narrative of Harish et al. 2,3]. A somehow similar table fact-checking can be found in our eLetter response [5].

## Their claim that our rooting approach uses outgroup taxa is incorrect

We do not *“use a hypothetical pseudo-outgroup, … an artificial all-zero taxon … to root the ToL*” [2] nor *“an ancestor that is assumed to be an empty set of protein domains”* as outgroup to *“create specific phylogenetic artifacts”* [3]. No outgroup taxon (presumably extant or artificial) was ever used/defined in our study [4], demonstrating they are confusing outgroups with ancestors (Table 1). We are therefore puzzled by the indirectly rooted ToLs they build in their attempt to mimic and undermine our methods [3], as well as the putative sensitivity to addition of new taxa, especially because these taxa represent organisms that engage in extreme forms of cellular endosymbiosis (please read below for the rationale for why then viruses were included). Even their tree searches were conducted differently, failing to minimize Farris’ *f*-values and being dependent on the location of the root. Contrary to their claims, our rooting method is grounded in early and well-established cladistic formalizations [7,8] and is *direct* because it polarizes character transformations with information solely present in the taxa being studied (the ingroup), distinguishing ancestral from derived character states and rooting the trees both empirically and *a posteriori*. In fact, and crucially, our rooted trees comply with Weston’s generality criterion [9,10], which states that as long as ancestral characters are preponderantly retained in descendants, ancestral character states will always be more general than their derivatives given their nested hierarchical distribution in rooted phylogenies. Biologically, domain structures spread by recruitment in evolution when genes duplicate and diversify, genomes rearrange, and genetic information is exchanged. This is a process of accumulation and retention of iterative homologies, such as serial homologues and paralogous genes [10], which is global, universal and largely unaffected by proteome size. This same process is widely used to generate rooted phylogenies from paralogous gene sequences. The Lundberg method [8], which does not attach outgroup taxa to the ingroup as they claim, simply enables rooting by the generality criterion [11].

## Their confusion of a priori and a posteriori character polarization questions their understanding of cladistic methodology

Their claim that we use the pseudo-outgroup to polarize character state changes *a priori* is inconsistent with our methodology of first reconstructing an undirected network and then polarizing character transformations with a direct method that complies with Weston’s rule [4]. They miss the fact that rooting is not a neutral procedure. While the length of the most parsimonious trees is unaffected by the position of the root, making *a priori* polarization unnecessary [7], rooting impacts the homology statements of the undirected networks [7,8]. *“The length of a tree is unaffected by the position of the root but is certainly not unaffected by the inclusion of a root”* [12].

## Their claim that small genome size affects rooting and induces attraction artifacts is conceptually and empirically false

Their recitation that organisms and viruses with small genomes (irrespective of their taxonomic affiliation) would be attracted to basal branches of our trees is incorrect. During searches of tree space and prior to rooting, we optimize character change in unrooted networks. This allows unrestricted gains and losses of domain occurrence or abundance throughout their branches. Thus, *rooting plays no role in defining unrooted tree topology* and cannot be distorted by genome size, which is a property of taxa (not individual characters changing in trees). This was already described in our supplementary text [4] and made explicit in a recent phylogenetic reconstruction study [13]. Polarization is only applied empirically *a posteriori*: *(a)* considering character spread in nested branches while accounting unproblematically for homoplasy (Weston’s rule), *(b)* searching for most parsimonious solutions with Lundberg while treating homologies as taxic hypotheses, and *(c)* allowing gradual and punctuated build-up of evolutionary emergence of protein structures, including gain and loss, that complies with the principle of spatiotemporal continuity (PC), Leibnitz’s *lex continui*. These three mutually supportive technical and biological axiomatic criteria were confirmed experimentally by Venn group distribution in ToDs and by visualizing clouds of proteomes in temporal space (*see* Figs. 5 and 8 of [4]). Felsenstein’s suggestion of inverse polarization [14] of our ordered (Wagner) characters, which can be polarized in only two directions, produced suboptimal trees (e.g., Figs. 3 and 4 of [6]). Empirically, plotting the node distance (*nd*) for each terminal node (i.e. taxa) from the root node of a ToL - on a scale from 0 (most basal) to 1 (most recent) - against the genome sizes of taxa showed substantial scatter (*rho* = 0.80), poor lineal fits (several peaks and troughs with 100 iterations of the LOWESS fitting method and a smoothing of *q* = 0.05), shallow monotonic increases (flat lines in Archaea and Bacteria), and no distortions/mixing of taxa among the four supergroups, viruses, Archaea, Bacteria and Eukarya. Figure 1A shows the plot for viral taxa located at the base of the rooted ToL. Genomes of similar sizes were scattered throughout the *nd* axis suggesting that genome size was not a significant determinant of taxa position in our trees. Similarly, genome size scatter for individual *nd* increased towards the base of the trees, including scatter for supergroups (*rho* values of 0.55, 0.63, 0.75 and 0.85 for viruses, Archaea, Bacteria and Eukarya respectively), debunking the alleged basal attraction artifact. The order of appearance of supergroups matched their proteomic complexity, from simple to complex, which also matched scaling patterns of use and reuse of structural domains in proteomes [6]. This emerging property of trees supports evolution’s PC.

**Figure 1.**
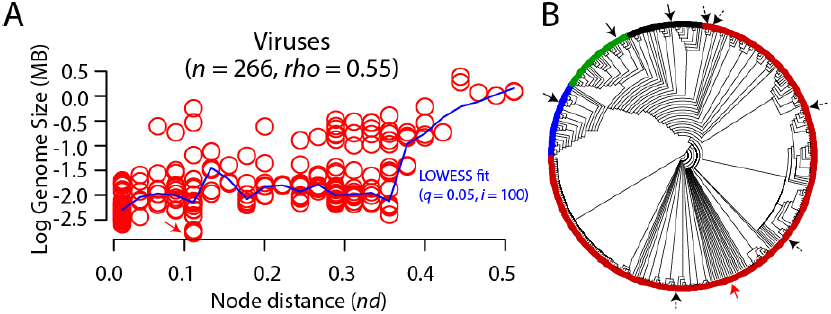
Trees of proteomes are robust and insensitive to the effects of genome size. **A**. Scatter plot describing the relationship between genome size and node distance (*nd*) for viral taxa used in our study [4]. For clarification, our data set included 266 viral proteomes including up to 5 viral representatives from each known viral family/order and belonging to each of the seven replicon types seen in viruses, as in [4]. The blue line describes the nature of the relationship, as determined by the Locally Weighted Regression Scatter Plot Smoothing (LOWESS) method, which obtains a smoothed curve by fitting successive regression functions. The red arrow indicates the smallest virus of the genomic set, the bat cyclovirus (1.7kb genome and encoding a single fold superfamily domain). High scatter values of the plot (*rho* = 0.55) indicate no ‘small genome attraction’ artifact that would pull small genomes toward the base of the tree, i.e. towards *nd* = 0. **(B)** The single most parsimonious tree (length = 45,935, retention index = 0.83, *g*_1_= −0.31) describing the evolution of 102 cellular organisms and 266 viruses (described in [4]). The smallest proteomes in each cellular supergroup are represented by black arrows (see text for description). The smallest viral proteome (bat cyclovirus) is labeled with a red arrow. Viruses sampled by Harish et al. [3] are indicated with dashed arrows. The names of taxa are not shown because they would not be visible. Instead, the positions of terminals were colored according to supergroup, green (Eukarya), blue (Bacteria), black (Archaea) and red (viruses).

Labeling the phylogenetic positions of the smallest proteomes in our trees confirmed that the smallest genomes were neither attracted towards the root nor caused distortions in the four-supergroup ToL (see also Fig. 7A in [4]). Here, Figure 1B showcases the phylogenetic positions of the “smallest” proteomes in each of the four supergroups (Archaea, Bacteria, Eukarya, and viruses) in our ToL. For clarification, the smallest proteomes included *Ignicoccus hospitalis* (Archaea, 213 fold superfamilies), *Lactobacillus delbrueckii* (Bacteria, 261), *Ashbya gossypii* (Eukarya, 326), and the bat cyclovirus (virus, 1). We also labeled the 9 viruses (4 RNA and 5 dsDNA) that produced phylogenetic distortions in the trees of Harish et al. [3]. Following the Harish et al. [3] logic, one should expect for the smallest proteomes to “fight” for the basal position within each of the four-supergroup subtrees or appear together closest to the root, irrespective of their taxonomic affiliation [2,3]. Contrary to that, the smallest proteomes appeared at well-derived positions (see arrow pointers) and did not cause any phylogenetic distortions or mixing of taxa from different supergroups! Even the smallest virus (the 1.7kb bat cyclovirus encoding a single fold superfamily) did not appear at the cluster of basal RNA viruses but was clustered with its closest evolutionary relative, the Dragonfly cyclovirus at more derived positions (red arrow in Figure 1B and at *nd* = 0.11 in Figure 1A). Thus, claims that *“the rooting in viral lineages is an inevitable consequence of pre-specifying ‘0’ or ‘absence’ as the ancestral state”*, that *“the position of the root depends on the smallest genome in the sample”* [2], and that we *“recognized anomalous effects of including small genomes in reconstructing the ToL”* [3] are therefore conceptually and empirically false. The contradictory results obtained by Harish et al. [2,3] therefore stem from (i) their misunderstanding of our rooting approach and cladistics, and (ii) intentionally/unintentionally missing the taxa sampling rationale that was well explained in our study [4]. To clarify, we took great care in sampling an equal number of cellular organisms from each supergroup (34 each) and at least five viruses from each family/order (87 viral families and 266 viruses were represented in our trees [4]). Indeed, our ToL (Figure 7 in [4]) revealed many monophyletic groups of viruses that were consistent with ICTV classifications and was far more complicated than showed/claimed by Harish et al. [3]. In turn, and in their haste to claim phylogenetic distortions, Harish et al. [3] likely handpicked problematic taxa (only 9 viruses in comparison to our 266) to claim phylogenetic distortions and to misguide the interpretation of our phylogenomic methodology (please also see Section 5). This was a crucial mistake since we had explicitly stated that taxa must be sampled broadly for the success of genome composition based phylogenies (see supplementary material in [4]). Even, the addition of the extremely reduced *Rickettsia prowazeki* and *Nanoarchaeum equitans* genomes (originally excluded because of their lifestyles) to our dataset, did not cause any topological distortions to our trees (data not shown), in contradiction to claims of Harish et al. [3].

## Confusion of characters and taxa bootstraps their preconceptions

In rushing their unsupported claim that the rooting of trees of structural domains (ToDs) is also unreliable and affected by small genome size, they wrongly considered ToDs as being *“uninterpretable in terms of the definition of the* (domain) *superfamilies which it comprises”*, because homology *“within”* superfamilies *“can be ascertained based on similarity of sequence, structure and function”*. But superfamilies are the taxa and proteomes the characters, and definitions of taxa (superfamily hidden Markov models) do not need to follow either definitions of characters (superfamily growth in proteomes) or statements of homology tested in ToDs. What is *‘uninterpretable’* however is the putative effect of genome size on ToDs, since each proteome embodies a character, which by definition (Kluge’s Auxiliary Principle) is independent of others. So fiction bootstraps their preconceptions, including the idea that domains, the evolutionary units of proteins, do not evolve. Are 1,200 structural folds fortuitous findings or the makings of intelligent cause? Where does significant evolutionary signal of the ToDs, including a match to the geological record [15], come from? Even an exploration of the mapping of functions in genotype space shows the centrality of structure in defining evolutionary constraints [16].

## Their analysis failed to avoid sampling pitfalls and problematic taxa

They confused exclusion of taxa engaged in obligate parasitism and symbiosis in cellular organisms with avoidance of genome size attraction artifacts, when in fact our intention was to exclude organisms with ill-defined hologenomes of holobiont collectives (the host and its associated organismal communities), which are known to complicate definitions of taxa [17,18]. No such attempt was extended to the viral supergroup since one hallmark of viruses is harboring a life cycle with strict dependence on a host. We explored their dataset and found that sampling of taxa was limited and imbalanced (Table 1) and included questionable taxa that were likely cherry-picked from extreme proteomic outliers (e.g. Fig. 1C in [4]), and sometimes selected outside our initial sampling (e.g. *Cand*. Nausia deltocephalinicola). For example, *Cand*. Tremblaya princeps included in their trees (Figure 2B in [3]) is part of a three-pronged endosymbiotic organismal system (bug in a bug in a bug) [19]. Its genome encodes only 55 universal domain superfamilies. It is not considered an independent organism since it depends on its host (*Planococcus citri*) and its endosymbiont (*Cand*. Moranella endobia) to synthesize essential metabolites [20]. To quote López-Madrigal et al. *“The genome sequence reveals that ‘Ca. Tremblaya princeps’ cannot be considered an independent organism but that the consortium with its gammaproteobacterial symbiotic associate represents a new composite living being”* [20]. Similarly, *Cand*. N. deltocephalinicola is an obligate endosymbiont of leafhoppers, which harbors the smallest known bacterial genome and encodes only 53 universal superfamilies [21]. These extreme proteomic outliers do not bias tree reconstructions because of their size nor induce *“grossly erroneous rootings”* [3]. Instead, their hologenomes arise from relatively modern genomic exchanges and recruitments likely resulting from complex trade-off relationships that complicate the dissection of their evolutionary origin and their definition as single taxon in phylogenetic data matrix. Phylogenetically, they represent ‘problematic’ taxa that should be excluded from analysis pending further understanding of their genetic make up. The intentional inclusion of problematic taxa is expected to generate biased reconstructions; see [22] for a dinosaur phylogeny example and the detection of problematic taxa with double decay analysis. Instead, our decision to exclude organisms that do not engage in free-living relationships avoids these kinds of pitfalls [23]. In contrast to the effect of lifestyles, comparing trees built from information-related and nonrelated domain families uncovered that tree reconstruction is refractory to biases resulting from horizontal gene transfer [23], a much more serious putative effect on tree reconstruction.

They state that our Venn diagrams and summary statistics of domain distributions in supergroups are unconvincing in light of other comparative genomic analyses [24,25]. However, Abroi and Gough [24] argued that viruses may be a source of new protein fold architectures, a conclusion strongly supported by our Venn analysis, and Abroi [25] showcases the distribution and sharing of domain superfamilies between viral replicon types and cells, which is remarkably consistent with our analysis [4]. There are no irreconcilable differences between these studies. Instead, our phylogenomic analysis dissects the alternative evolutionary scenarios that can be posed with the comparative method [4,25]. For example, we also mapped virus-host information to the Venn distribution of protein fold superfamilies and identified 68 fold superfamilies that were (remarkably) shared by archaeo-, bacterio-, and eukaryoviruses (Fig. 3 in [4]). These superfamilies included many ancient folds involved in cellular metabolism and hinted towards an origin of viruses predating the origin of modern cells. Harish et al. [3] therefore ignore the plethora of evidence given in our article (Figs. 1-8 in [4]) supporting the ancient origin of viruses and instead rely on customized phylogenetic trees to claim invalidity of our results.

### Conclusions

Harish et al. [2,3] believe we root our ToLs *a priori* with an indirect method and an outgroup taxon they confuse with an ancestor, when in fact we root our ToLs *a posteriori* using a direct method compliant both with Weston’s rule and experimentation. They claim our rooting method attracts organisms and viruses with small genomes to the base of rooted trees when in fact tree topology is established during unrooted tree optimization and prior to character polarization. It is ironic that attempts to controvert our direct methods of character polarization come from authors that are themselves proponents of the use of polarized characters, but with arbitrary transformation costs carefully engineered *a priori* to attract large eukaryotic genomes to the base of their trees [1]. These characters violate the ‘triangle inequality’ [26], a fundamental property conferring metricity to phylogenetic distances. Its violation invalidates phylogenetic reconstruction [27]. Quoting the cladistics opinion of W. C. Wheeler [26] on the validity of characters with arbitrary transformation costs: *“Although the matrix character optimization algorithm does not require metricity, biologically odd results may occur otherwise. As an example, an additional state could be added to an existing set, with very low transformation cost to all other elements (σ_k,0_ < ½ min σ_i,j_). The median state at all internal 2 vertices (V \ L) would then be this new state for all trees, no matter what the leaf conditions were”* (page 184). He further illustrates the unforeseen consequences of optimization with non-metric distances using the well-known NP-hard ‘traveling salesman problem’, a salesman that wishes to visit a number of cities while minimizing travel time. This is a notorious task known to require considerable travelling optimization. The use of non-metric distances however makes a city have zero distance to all other cities, creating a ‘wormhole’ in space-time that allows to reach all cities at zero cost. Such bizarre property has dire consequences for the recovery of a correct tree and invalidates the Harish et al. [1] approach. Add to this the fact that their loss-favoring step matrices require that they be solely optimized on a rooted tree, making rooted tree reconstruction prone to ‘large genome attraction’ artifacts. In addition to this self-inconsistency, transformation costs also violate genomic scaling and processes responsible for scale-free behavior of proteins, challenging evolution’s PC and artificially forcing biological innovations to the base of universal trees [6], effectively resulting in an ancestral clumping of irreducible complexity.

